# Liver X Receptor activation regulates genes involved in lipid homeostasis in developing chondrocytes

**DOI:** 10.1101/705269

**Authors:** Margaret Man-Ger Sun, Frank Beier

## Abstract

**Objective:** Osteoarthritis (OA) is the most common type of arthritis and causes debilitating symptoms and decreased quality of life. Currently available treatment options target symptoms but do not address the underlying issue of joint tissue degeneration. As such, a better understanding of the molecular mechanisms maintaining cartilage health is needed for developing novel therapeutic strategies. Liver X Receptors (LXRs) are nuclear receptors that have been previously shown to offer protection against OA. This is potentially due to suppression of chondrocyte hypertrophy in endochondral bone growth in response to LXR activation. In order to better understand the regulatory mechanisms behind this effect, we aimed to systematically examine LXR’s effects on growth plate chondrocyte gene expression.

**Methods:** Primary chondrocytes isolated from the long bones of E15.5 mice were treated with the specific LXR agonist, GW3965, and RNA was isolated for Affymetrix microarrays followed by real time qPCR validation. Bioinformatics analyses were performed using Gene Ontology (GO) and KEGG pathway analysis. Immunohistochemistry was conducted to examine protein localization of LXR and identified targets in GW3965-treated E15.5 tibiae compared to control.

**Results:** Activation of LXR in primary growth plate chondrocytes resulted in differential regulations of various genes involved in lipid metabolism, including several genes involved in cholesterol efflux. This pattern was compared to LXR activation in immature murine articular chondrocytes (IMACs), which revealed similar roles in lipid homeostasis. Immunohistochemical analysis of LXR and its identified targets Abca1 and Srebf1 revealed preferential protein localization to pre-hypertrophic and resting chondrocytes in GW3965-treated tibial growth plates compared to controls.

**Conclusion:** Our findings show for the first time that LXR activation alters expression of lipid metabolism genes in growth plate chondrocytes, in part through activation of molecules responsible for cellular cholesterol efflux. This provides insight into potential mechanisms through which LXR regulates cellular metabolism to alter chondrocyte behavior and phenotype.

## Introduction

Osteoarthritis (OA) is a prevalent degenerative joint disease that globally affects 10% of men and 18% of women over the age of 60 (1,2). Currently available treatment options merely alleviate symptoms and attempt to improve function through approaches such as joint replacement surgery, but do not address the underlying cause (3). Thus, there is an urgent need for a better understanding of the cellular and molecular mechanisms that contribute to the pathogenesis of OA. Osteoarthritis is a multifactorial disease with a complex causative and progressive process. The contributing sources to the pathophysiology of primary OA include age, genetics, gender, weight and metabolism, whereas secondary OA is primarily attributed to joint injury (4,5). Although OA may affect tissues of the entire joint, the characterizing feature of OA is the progressive loss of the structure and function of the articular cartilage (5,6). Cartilage degradation in OA is generally due to the loss of the balance between synthesis and degradation of the extracellular matrix (ECM) responsible for maintaining cartilage homeostasis and function, and changes in chondrocyte phenotype also contribute to this pathology (5,7,8).

Certain cellular and molecular events in endochondral bone formation have been found to be recapitulated in OA pathogenesis, where disruptions of chondrogenic differentiation can lead to the onset of OA (9). In particular, ectopic chondrocyte hypertrophy within the articular cartilage has been shown to act as an OA-promoting factor (9–11). This is demonstrated by protection against injury-induced OA in mice lacking drivers of chondrocyte hypertrophy such as Runx2 or Hif2α (12–14). It remains unclear whether it is chondrocyte hypertrophy itself, or simply the induction of hypertrophy-associated genes, or the combination of both that contributes to cartilage degeneration in OA (15). Nevertheless, the differentiation and phenotypic changes that developing growth plate chondrocytes undergo are usually resisted in healthy articular chondrocytes, allowing for a stable and permanent structure for joint articulation (16). In OA, this resistance is lost, resulting in development of hypertrophic features in articular chondrocytes. Better understanding of the regulatory mechanisms that control the phenotypic disparity between growth plate and articular chondrocytes can help shed light on the cellular changes that initiate this transition.

Nuclear receptors are a class of transcription factors activated by small lipophilic substances to regulate numerous developmental and homeostatic processes (17). Previous studies from our laboratory and others have identified nuclear receptors to play important roles in cartilage biology and osteoarthritis (18–23). Most notably, Liver X Receptor (LXR) activation appears to protect from OA (21). Our work has demonstrated that activation of LXR regulates various lipid metabolic processes within post-natal immature murine articular chondrocytes (IMACs) (19). We further showed LXR activation to suppress chondrocyte hypertrophy during endochondral bone growth, formulating a novel potential underlying mechanism of LXR’s protection against OA (23). However, the signaling pathways and target genes mediating this suppression remain to be elucidated.

The objective of this study was to identify target genes of LXR in developing growth plate chondrocytes. We performed microarray analyses on RNA extracted from primary chondrocytes isolated from embryonic day 15.5 (E15.5) mouse long bones treated with or without an LXR agonist. Differentially regulated genes were subsequently validated at both the mRNA and protein level, where we show changes in gene expression after LXR agonist treatment to primarily affect lipid metabolism. We also show for the first time LXR’s ability to regulate target protein localization within the developing growth plate. These results suggest LXR’s regulation of chondrocyte differentiation may be due to their role in altering lipid metabolism in growth plate chondrocytes.

## Methods

### Animals and materials

All procedures involving mice were performed in accordance to the Canadian Council on Animal Care (CCAC), and followed the animal care and use protocol approved by the Council on Animal Care at Western University (Canada). Timed pregnant CD1 mice were obtained from Charles River Laboratories. The LXR agonist, GW3965, and all real-time PCR reagents and primers were obtained from Sigma-Aldrich. All organ culture and cell culture media components were obtained from Sigma-Aldrich and Thermo Fisher Scientific, unless otherwise stated.

### Micromass culture

E11.5 limb buds were isolated from CD1 mouse embryos and segmented, dissociated, and cells were cultured in micromass cultures, as previously described (23). Micromass cultures were treated with either dimethylsulfoxide (DMSO) as vehicle control, or varying concentrations of GW3965 (1 μM or 5 μM), and cultured for a period of 12 days with media and treatment replenished every 24 hours.

### Primary chondrocyte culture

Primary chondrocytes were isolated from long bones of E15.5 CD1 mice (femur, tibia, radius, ulna, fibula, humerus, scapula) and cultured as previously described with modifications (24). To remove muscle and other associated tissue, isolated long bones were subjected to a 15 minute incubation in 500 μL of trypsin at 37°C under agitation. This was followed by a 1.5-2 hour incubation in Collagenase-P (3 mg/mL, Roche) diluted in Dulbecco’s Modified Eagles Medium (DMEM) supplemented with 10% fetal bovine serum (FBS) at 37°C under 5% CO_2_. Digested cells were strained with a 40 μm cell strainer, then plated on 1.2% agarose. Cells were allowed to recover overnight before commencement of treatment using DMSO as vehicle control, or GW3965 at 1 μM concentration for a period of 24 hours.

### RNA isolation and microarray analyses

Following treatment, RNA was isolated from primary chondrocytes using TRIzol™ (Invitrogen) as per manufacturer’s instructions and as previously described (19). In brief, cell homogenization with TRIzol™ was followed by the addition of chloroform for phase separation and the upper aqueous phase was removed. RNA was precipitated using 100% isopropanol and washed with 70% ethanol. RNA was allowed to air-dry prior to resuspension in RNAse-free water, then quantified using a Nanodrop 2000 spectrophotometer (Thermo Fisher Scientific). RNA integrity was determined using the Agilent 2100 Bioanalyzer, and samples with RNA integrity number (RIN) greater than 8 were used for microarray analyses at the London Regional Genomics Centre, Robarts Institute (London, ON, Canada).

Total RNA (200 ng/uL) from three independent cell culture experiments underwent 2 rounds of amplifications, then labeling and hybridization to Affymetrix GeneChip™ Mouse Gene 2.0 ST Array (Thermo Fisher Scientific) containing probe sets representing > 28,000 coding transcripts, as previously described (19). The complete microarray data set will be deposited at the publicly accessible Gene Expression Omnibus (GEO) database, and NCBI accession number provided.

### Bioinformatics analyses

Microarray data was analyzed using the Partek Genomics Suite v6.6, where probe data was normalized and annotated, and ANOVA conducted to identify differentially expressed genes between control and treated conditions. Genes with p<0.05 and a minimum of 1.5-fold change were considered significantly changed and used for subsequent gene list creation and bioinformatics analyses. Gene Ontology (GO) and Kyoto Encyclopedia of Genes and Genomes (KEGG) pathway analysis was performed to generate GO term and pathway enrichment. Data was prioritized based on enrichment score (negative natural log of the enrichment p-value) to indicate hierarchy of level of gene representation in GO terms and KEGG pathways.

### Real-time quantitative polymerase chain reaction (RT-qPCR)

A subset of RNA isolated from E15.5 primary chondrocytes was used for gene expression analysis through real-time quantitative PCR, as previously described (19,23,25). Complimentary DNA (cDNA) synthesis was conducted using the iScript™ cDNA Synthesis Kit with 500 ng of starting RNA (Bio-Rad Laboratories), and combined with 50 μM of forward and reverse primers (for primer sequences, please see Supplementary Table 1) as well as iQ™ SYBR^®^ Green Supermix (Bio-Rad Laboratories). Expression values were normalized to the internal control Beta-actin (*Actb*) or Glyceraldehyde 3-phosphate dehydrogenase (*Gapdh)*, calculated using the ΔΔCT method, and expressed relative to control cells treated with DMSO.

### Tibiae organ culture and immunohistochemistry

Tibiae were microdissected from E15.5 CD1 mice and cultured for a period of 6 days in the presence of DMSO as vehicle control, or 1 μM of GW3965 as treatment as described (23). Media and treatment with GW3965 were replenished every 48 hours. At the end of the 6-day incubation period, tibiae were rinsed with phosphate-buffered saline (PBS) and fixed overnight with 4% paraformaldehyde (PFA) (Sigma-Aldrich), then embedded and sectioned at the Molecular Pathology Core Facility, Robarts Institute (London, ON, Canada). Immunohistochemistry was conducted as described with minor modifications (19,23). Paraffin sections were dewaxed and carried through a series of rehydration steps, followed by incubation in 3% H_2_O_2_. Antigen retrieval was performed with 0.1% Triton X-100 for 11 minutes, then slides were blocked with 5% goat serum in PBS, and incubated overnight at 4°C with primary antibodies against LXRα/β (sc-377260, Santa Cruz Biotechnology), Abca1 (PA1-16789, Thermo Fischer Scientific), or Srebp1 (PA1-337, Thermo Fischer Scientific). Secondary antibody incubation was followed by colorimetric detection with diaminobenzidine (DAB) substrate-chromogen solution (Dako Omnis, Agilent) and counterstaining with methyl green solution.

### Statistical Analysis

All data were collected from at least three independent experimental trials. Data were represented as mean ± 95% confidence intervals (CI), and p-values less than 0.05 were considered significant. Statistical analysis was performed using a parametric, unpaired two-tailed Student’s *t*-test using GraphPad Prism 6.0.

## Results

### Differential gene expression in response to LXR agonism in growth plate chondrocytes

Our laboratory previously reported LXR activation through GW3965 treatment to result in delayed chondrocyte hypertrophy in endochondral bone growth (23). In order to further understand the regulatory role of LXR in chondrocyte hypertrophy and growth plate development, we performed microarray analyses for differential gene expression in E15 primary chondrocytes treated with or without the LXR specific agonist, GW3965, for a period of 24 hours. Expression of 30 annotated genes was found to be significantly changed in response to LXR activation, with 7 genes down-regulated and 23 genes up-regulated (Supplementary Table 2).

Genes upregulated most highly with LXR activation include those encoding ATP-binding cassette sub-family G member-1 (*Abcg1*), ATP-binding cassette sub-family A member-1 (*Abca1*), sterol regulatory element-binding transcription factor 1 (*Srebf1/Srebp1*), Lysophosphatidylcholine Acyltransferase 3 (*Lpcat3)*, Stearoyl-CoA desaturase (*Scd1*), and Acyl-CoA synthetase (*Acsl3*) [Fig. 1(a)]. Induction of these most highly changed genes was subsequently validated through quantitative real-time PCR [Fig. 2]. Real-time results revealed consistent up-regulation of target gene expression with GW3965 treatment, which is in-line with our microarray results.

**Fig. 1.**
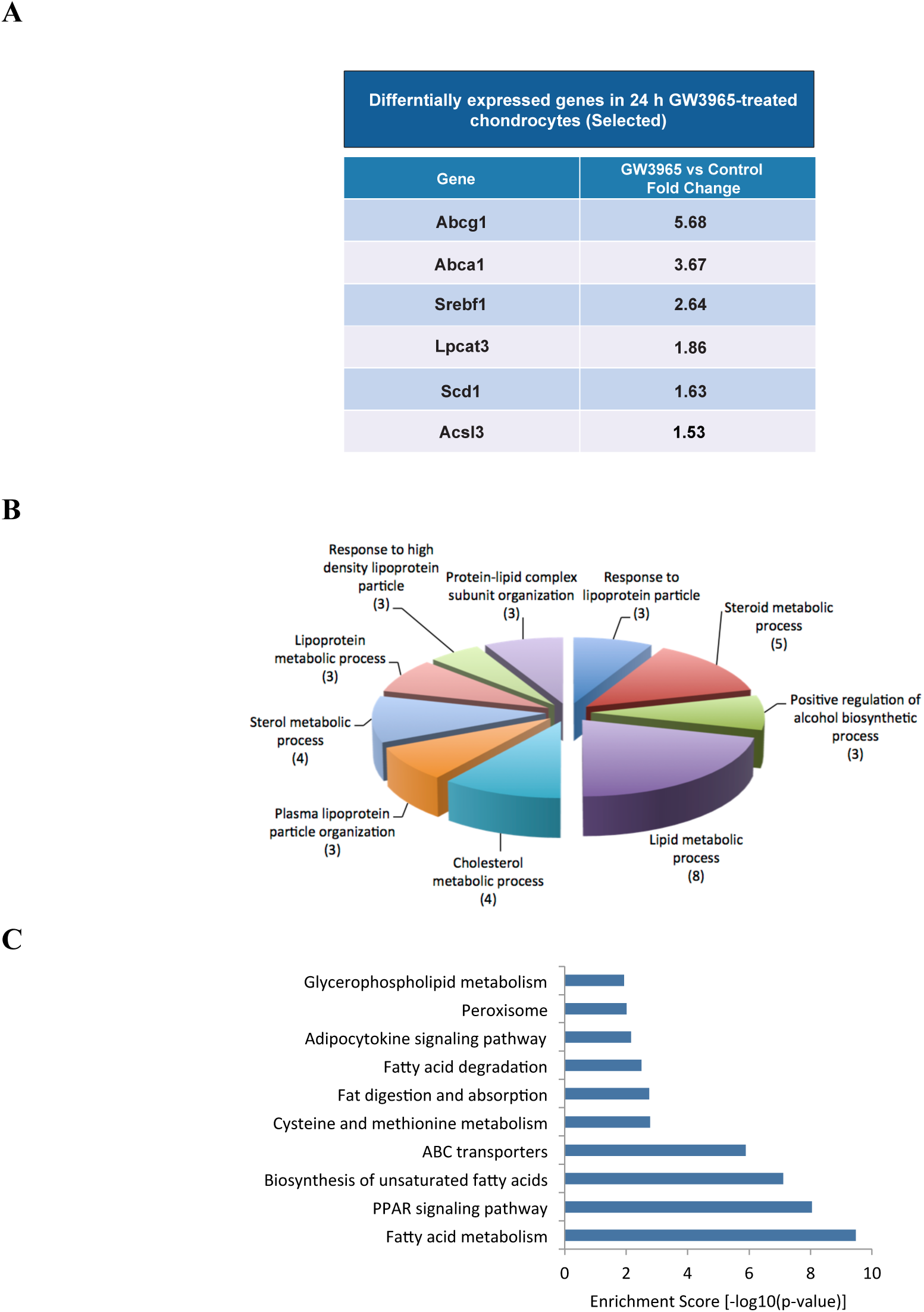
Microarray analyses of differential gene expression in GW3965-treated E15 primary chondrocytes. (A) RNA was isolated from E15 primary chondrocytes treated with either DMSO or 1 μM GW3965 for 24 hours and analyzed by Affymetrix microarray. N=3. (B) Gene Ontology (GO) groups were categorized using Partek and prioritized by enrichment scores. The top 10 groups and the number of regulated genes included in each GO group are shown. (C) KEGG pathway annotations prioritized according to enrichment scores.

**Fig. 2.**
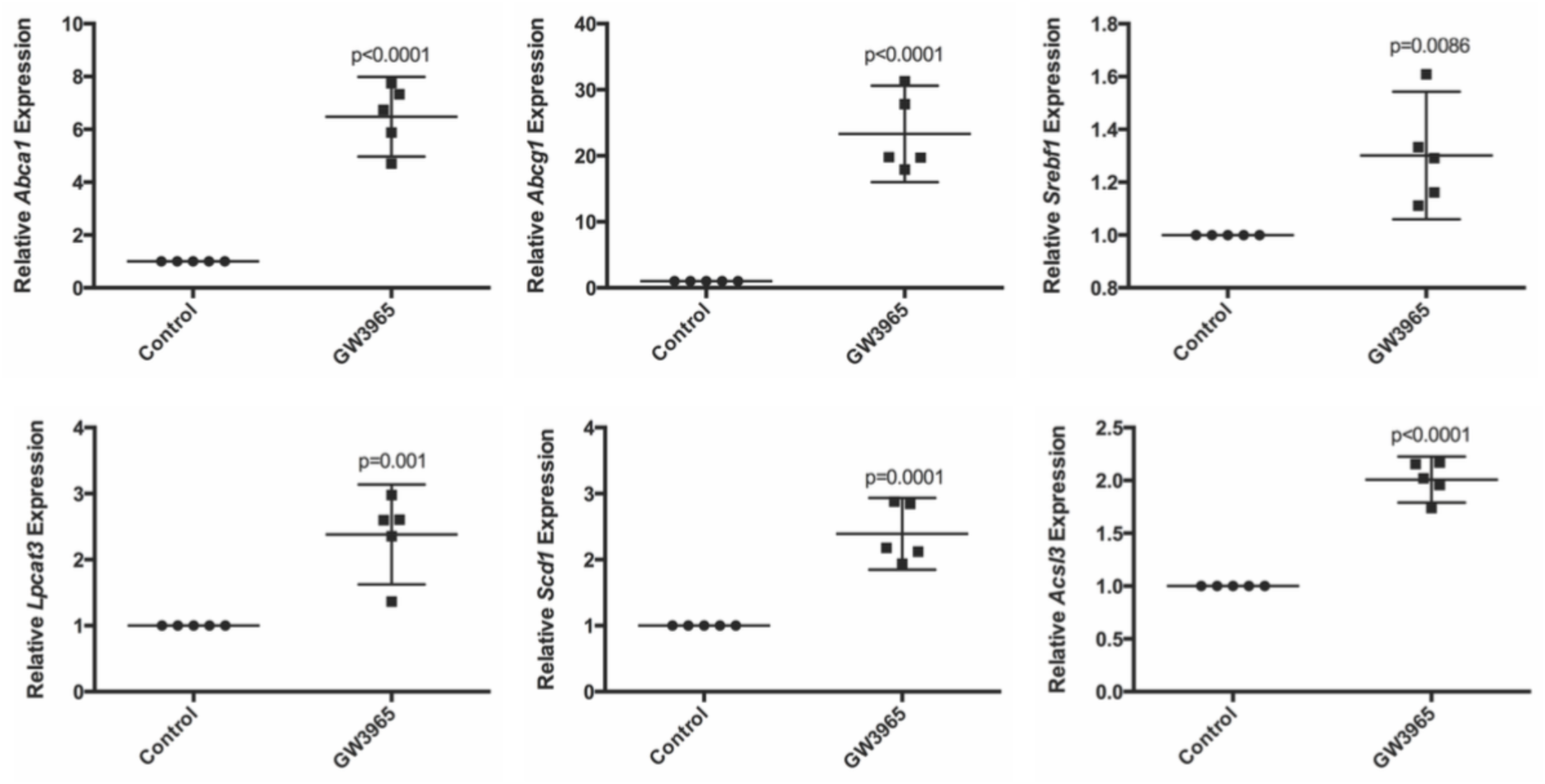
Real-time validation of differentially expressed genes in GW3965-treated E15 primary chondrocytes. RNA was isolated from E15 primary chondrocytes treated with either DMSO or 1 μM GW3965 for 24 hours, and quantitative real-time PCR was performed. The six most significantly up-regulated genes (selected) identified from the microarray, *Abca1, Abcg1, Srebf1, Lpcat3, Scd1*, and *Acsl3*, all demonstrated similar elevation in gene expression through quantitative real-time PCR. N=5, mean ± 95% CI.

In order to gain a better understanding of the potential roles of these differentially regulated genes, we conducted Gene Ontology and KEGG pathway analyses. This allows for the identification of represented cellular components, molecular functions, and biological processes, as well as associated pathways. The top 10 gene ontology groups prioritized by enrichment scores are primarily biological processes involved in lipid metabolism, including cholesterol metabolic process, lipoprotein metabolic process, and plasma lipoprotein particle organization [Fig. 1(b)]. Similarly, the top 10 KEGG pathways prioritized by enrichment scores indicate differentially regulated genes to be represented in various pathways of lipid homeostasis, including fatty acid metabolism, peroxisome proliferator-activated receptor (PPAR) signaling pathway, and adipocytokine signaling pathway [Fig. 1(c)]. Collectively, these analyses suggest the primary effect of LXR activation in growth plate chondrocytes to be control of cellular lipid homeostasis.

### LXR activation in growth plate and articular chondrocytes regulate similar lipid metabolic processes

To examine whether LXR activation through GW3965 results in similar gene expression patterns in growth plate chondrocytes and immature murine articular chondrocytes, we compared differentially expressed genes in each culture system by comparing our data sets described here to our previously published microarray data (19). Of the 30 up or down-regulated genes in GW3965-treated E15 chondrocytes, nine were found to be shared with genes regulated by GW3965 in the IMAC cultures [Fig. 3(a)]. The nine commonly changed genes are among the most up-regulated in both culture systems, and are mainly involved in lipid metabolic processes as well as cholesterol transport [Fig. 3(b)]. However, of the 133 genes differentially regulated by LXR activation in IMAC cultures, 124 identified genes were unique to this cell-type [Fig. 3(a)]. These data suggest LXR to exert similar lipid homeostatic functions in both embryonic and post-natal chondrocytes, while also regulating molecular functions unique to the phenotype of chondrocytes.

**Fig. 3.**
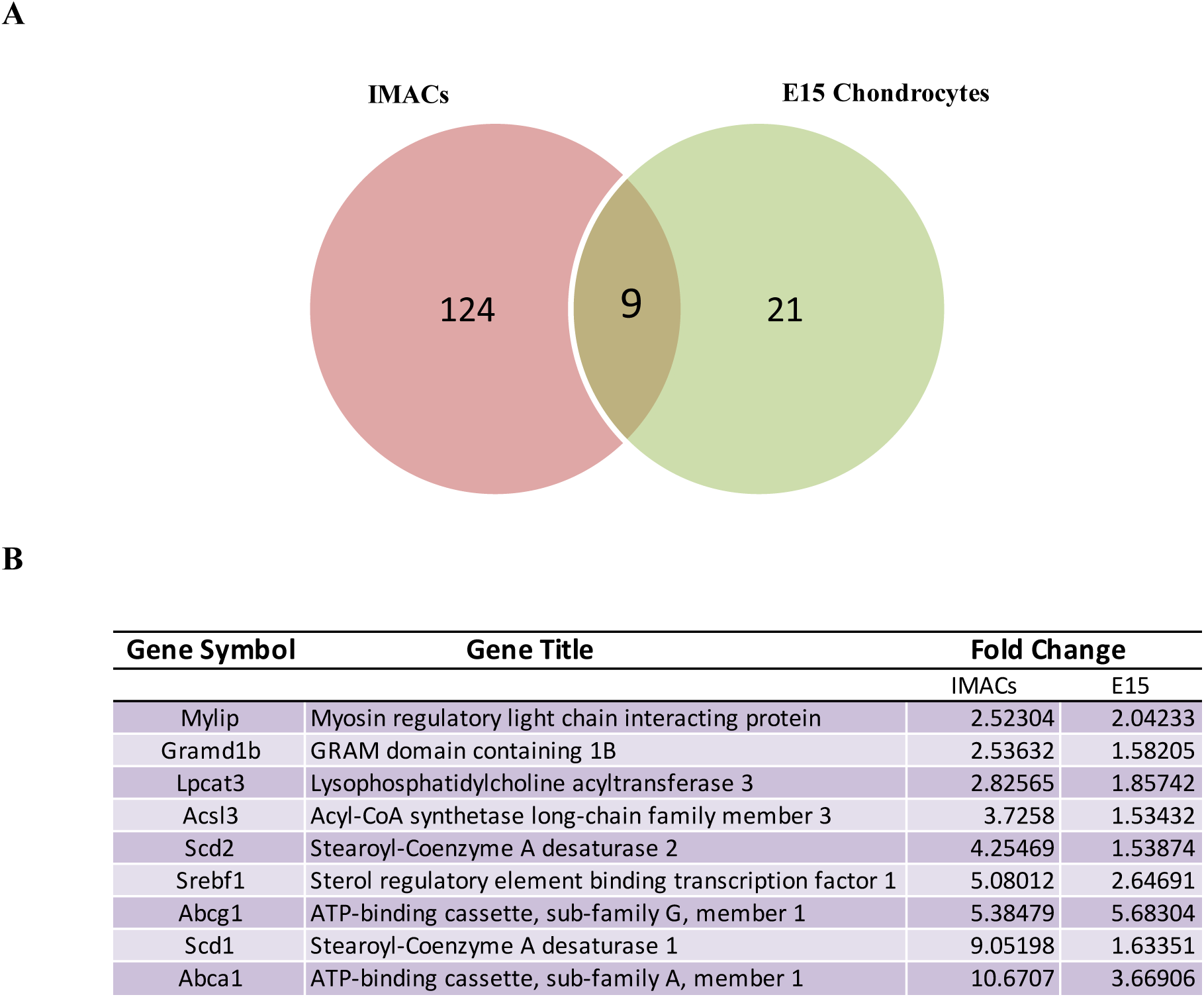
Comparison of differentially expressed genes in GW3965-treated E15 primary chondrocytes and GW3965-treated IMAC cultures. Genes up or down-regulated in GW3965-treated IMAC cultures were compared to genes up or down-regulated in GW3965-treated E15 primary chondrocytes. (A) Of the 133 genes differentially expressed in the IMAC cultures, 9 were also differentially expressed in the E15 primary chondrocytes. N=3. (B) The 9 common gene symbols, gene titles and their respective fold-changes are shown.

### LXR activation alters LXR gene expression and protein localization in developing chondrocytes

To gain insight into the effect of LXR activation on protein localization of LXR within the growth plate, immunohistochemical analysis was performed on E15.5 tibiae cultured in the presence of GW3965. Interestingly, despite being expressed throughout the growth plate in the control tibia, positive staining for LXR appeared to be more concentrated to the pre-hypertrophic and resting zones with GW3965 treatment [Fig. 4(a)]. Closer examination at 40x magnification revealed positive staining for LXR to be localized to the nucleus. Additionally, examination of LXR gene expression in chondrocytes undergoing hypertrophic differentiation in micromass cultures show significantly elevated expression of LXRα, and to a lesser extent LXRβ, at days 9 and 12 of culture [Fig. 4(b)]. Moreover, GW3965 appeared to increase LXRα and LXRβ transcript levels at later stages of micromass culture. These immunohistochemical and gene expression results suggest LXR activation to exert self-regulatory mechanisms based on chondrocyte differentiation phenotype, in particular for LXRα in pre-hypertrophic and hypertrophic chondrocytes.

**Fig. 4.**
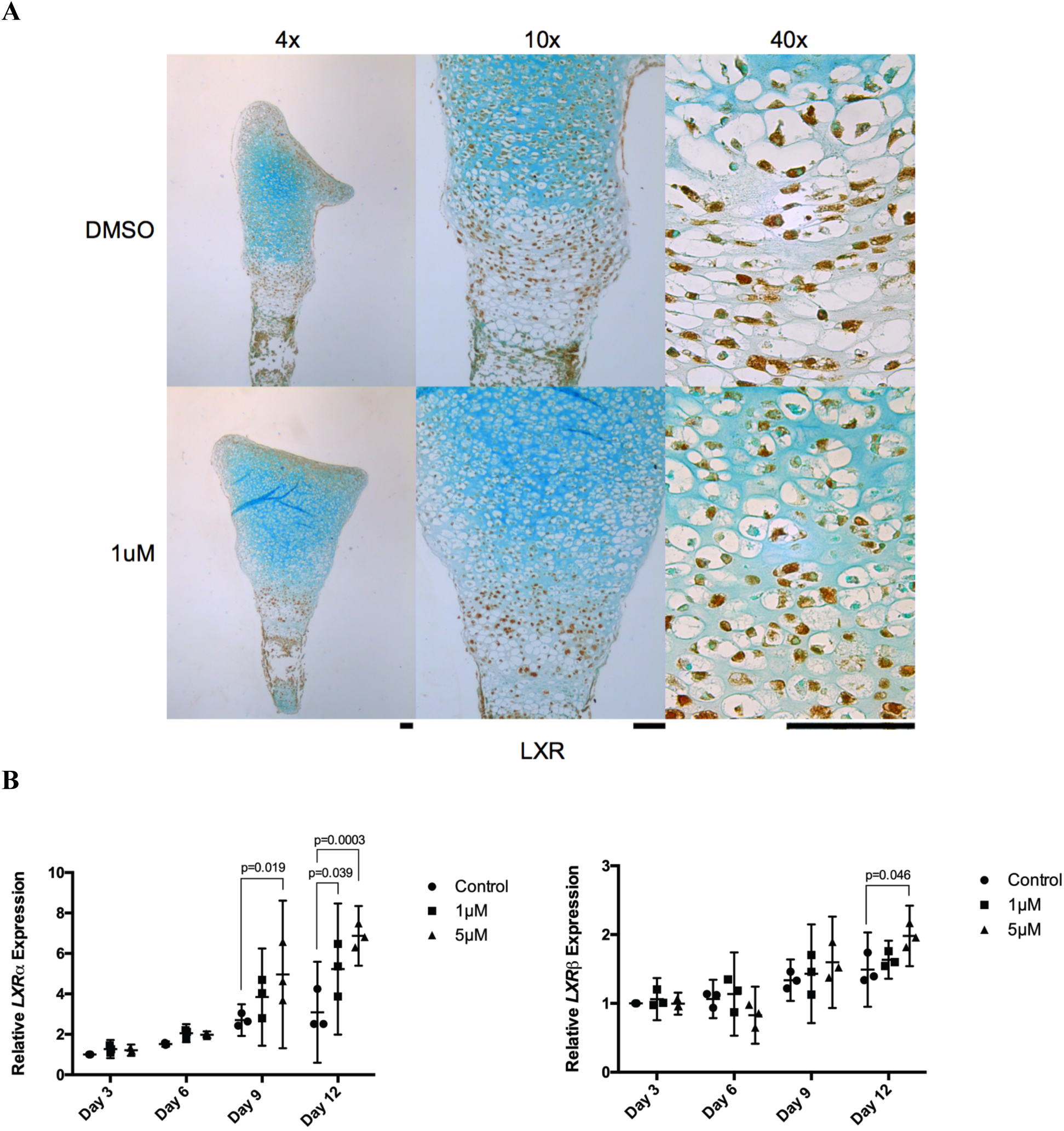
Effect of GW3965-treatment on LXR gene expression and protein localization. (A) Mouse tibiae were dissected at E15.5, grown in culture for 6 days and treated every 48 hours with DMSO or 1 μM of GW3965, then fixed and prepared for histology. Immunohistochemical analyses were conducted for LXR. Representative images show positive staining of LXR to be more concentrated to the resting and per-hypertrophic zones of GW3965-treated growth plates compared to control. Subcellular localization of LXR appears mainly nuclear. N=4, scale bar = 100μm. (B) RNA was extracted from micromass cultures treated with GW3965 or DMSO for a period of 12 days, and LXRα and LXRβ gene expression was examined through quantitative real-time PCR. N=3 - 4, mean ± 95% CI.

### LXR activation alters expression and localization of lipid metabolic targets in developing chondrocytes

We next conducted validation of differentially regulated genes upon LXR activation at the protein level through immunohistochemical analysis of GW3965-treated tibiae. For both Abca1 and Srebp1, protein localization appears to become more restricted to the pre-hypertrophic and resting zones upon LXR activation, mirroring what was observed with LXR protein expression [Fig. 5]. Positive staining of Abca1 was observed to be localized to the nucleus, the perinuclear area, and the cytoplasm [Fig 5(a)]. In contrast, subcellular localization of Srebp1 appears to be mainly nuclear [Fig. 5(b)]. Examination of differentially regulated lipid metabolic genes showed *Abca1, Abcg1, Scd1, Lpcat3*, and *Acsl3* to be significantly up-regulated by GW3965 throughout chondrocyte differentiation in micromass culture, but most markedly so during terminal chondrocyte differentiation [Fig. 6]. While *Srebp1* expression demonstrated a similar trend, its upregulation in response to LXR activation did not reach statistical significance in this culture system. Taken together, these findings suggest lipid metabolic targets of LXR to be regulated in a similar manner as LXR upon LXR activation, where their expression appears to be highly increased in pre-hypertrophic and hypertrophic chondrocytes.

**Fig. 5.**
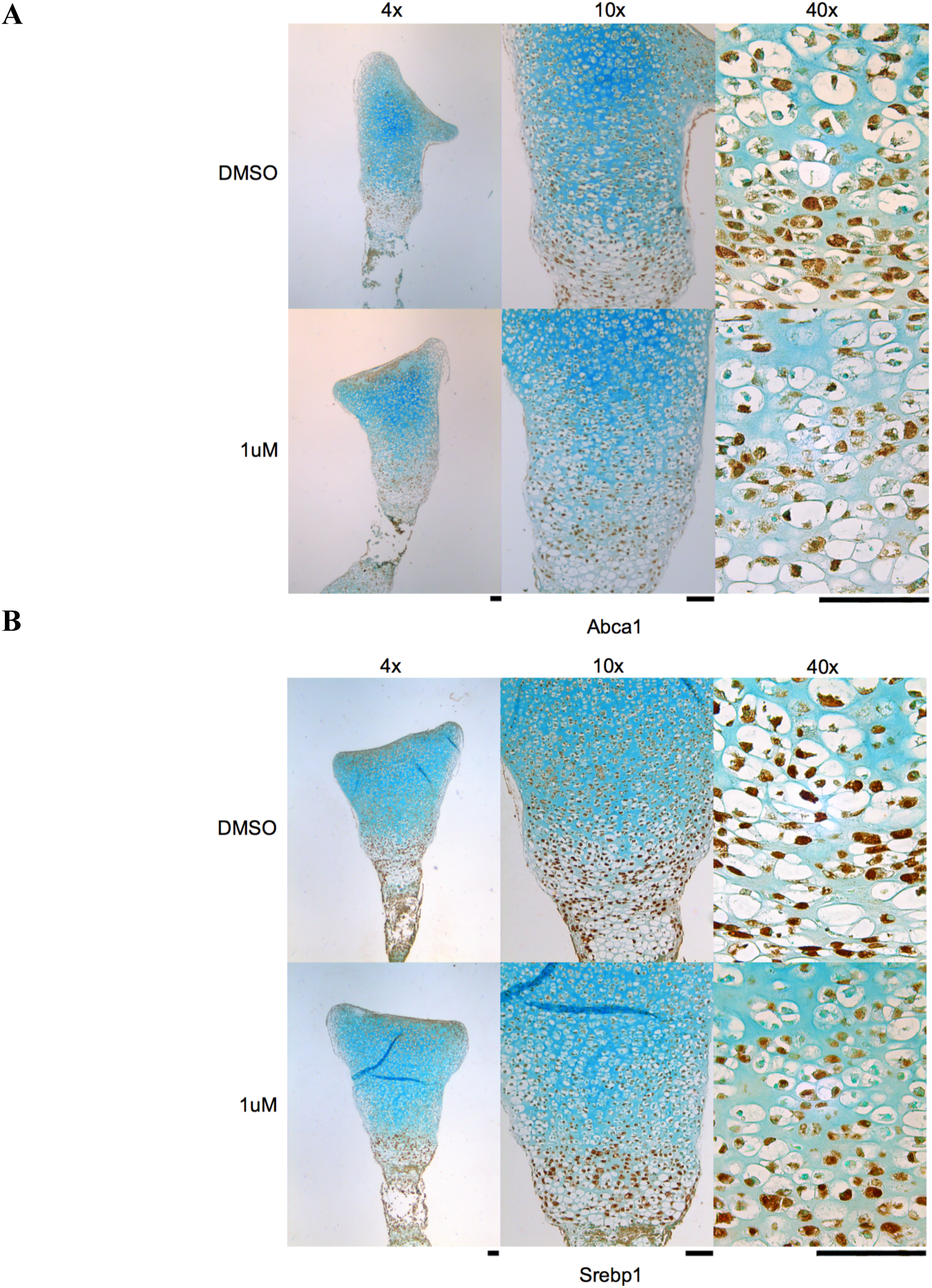
Effect of LXR activation on Abca1 and Srebp1 localization in the growth plate. Mouse tibiae were dissected at E15.5, grown in culture for 6 days and treated every 48 hours with DMSO or 1 μM of GW3965, then fixed and prepared for histology. Immunohistochemical analyses were conducted for Abca1 and Srebp1. Representative images show positive staining of Abca1 and Srebp1 to be concentrated to the resting and per-hypertrophic zones in GW3965-treated growth plates compared to control. Subcellular localization of Srebp1 appears to be mainly nuclear, whereas subcellular localization of Abca1 appears to be nuclear, perinuclear, and cytoplasmic N=4, scale bar = 100μm.

**Fig. 6.**
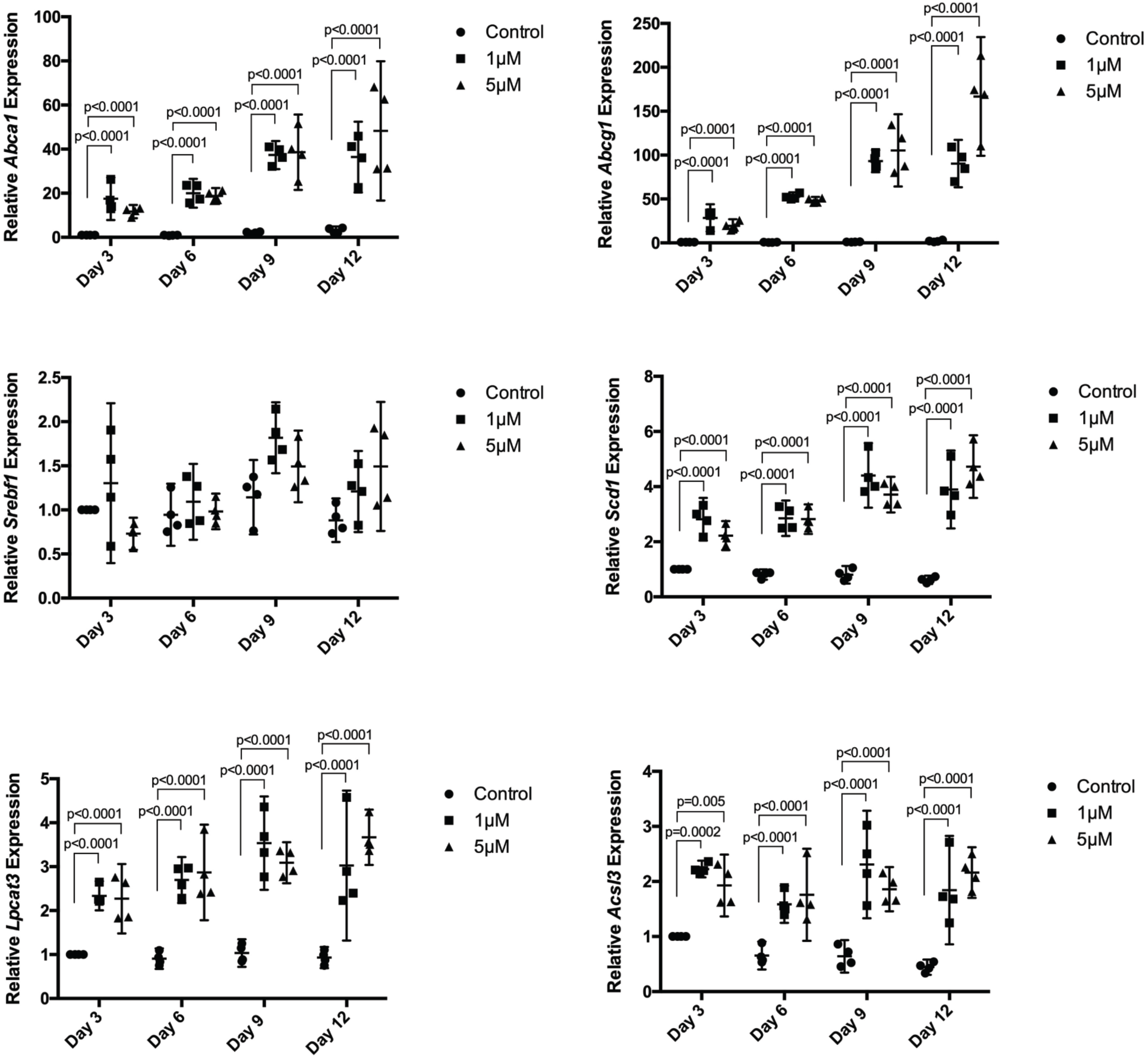
Effect of LXR activation on lipid metabolic gene expression in micromass cultures. Mouse embryonic limb buds were dissected at E11.5 and cells were cultured as micromasses for 12 days in the presence of DMSO, or 1 μM or 5 μM of GW3965. Gene expression was analyzed by quantitative real-time PCR and normalized to *Gapdh*. Expression of *Abca1, Abcg1, Scd1, Lpcat3*, and *Acsl3* was significantly increased with GW3965 treatment throughout the culture period, especially at days 9 and 12. Similar trends of increased expression was observed for *Srebp1*. N=4, mean ± 95% CI.

## Discussion

Previous studies have demonstrated LXR to play key functions in coordinating whole-body lipid homeostasis through highly metabolically active tissues such as the liver (26,27). However, much less is known regarding its roles in regulating lipid metabolism in cartilage. LXR’s potential therapeutic implications in OA has been clearly demonstrated by its chondroprotective effects in cartilage and in suppressing hypertrophic differentiation of developing chondrocytes (21,23); however, the underlying mechanisms behind these effects remain unknown. In the present study, we performed microarray analysis to examine transcriptional regulation by LXR in growth plate chondrocytes, and show for the first time that LXR activation regulates expression of genes involved in lipid homeostasis in differentiating growth plate chondrocytes.

Our microarray analysis revealed several essential proteins of reverse cholesterol transport to be amongst the most up-regulated genes identified in response to LXR activation. *Abca1* and *Abcg1* encode members of the ABC family of membrane transporters that promote transfer of cellular cholesterol to acceptor proteins in peripheral cells such as macrophages (28,29). *Abca1* and *Abcg1* are direct targets of LXR in human macrophages and keratinocytes (30–33), and can be induced by treatment of synthetic LXR ligands to result in decreased atherosclerosis pathology stemming from excessive cholesterol accumulation (32,34–36). Indeed, imaging mass spectrometry has shown cholesterol accumulation in the superficial area of human osteoarthritic cartilage (37), and cholesterol efflux gene expression is decreased in human osteoarthritic chondrocytes (38). Moreover, there is increasing evidence that intracellular cholesterol accelerates OA disease progression (39,40). Given our previous and current findings regarding LXR’s role in developing chondrocytes, it is possible that LXR’s protective effect in OA is mediated in part by their ability to increase cholesterol efflux and suppress chondrocyte hypertrophy (20,23). Our microarray results suggest LXR activation to exert similar lipid metabolic regulation in both growth plate and articular chondrocytes, and several studies have functionally linked this genetic regulation with changes in chondrocyte phenotype. Specifically, previous work from our lab has shown cholesterol administration to result in increased bone growth and chondrocyte enlargement similar to hypertrophy (20). Additionally, statin treatment and subsequent lowering of cholesterol has been shown to reduce protease activity and increase anabolic effects in chondrocytes, and was able to rescue chondrodysplasia phenotypes induced by FGFR3 mutation (39,41).

Similar to *Abca1*, our findings demonstrate LXR activation to result in increased *Srebp1* gene expression with protein localization mirroring LXR within the growth plate. Srebp1 is a transcription factor known for its role in regulating lipogenic enzymes such as Scd1, the rate limiting enzyme in the synthesis of monounsaturated fatty acids from saturated fatty acids (42). LXR’s ability to up-regulate Srebp1 may seem counterintuitive given the latter’s lipogenic nature, however studies have shown oleoyl-CoA to be an essential substrate for cholesterol esterification (43). Since Srebp1-directed induction of Scd1 is required for the synthesis of oleoyl-CoA, it appears that the LXR-Srebp1-Scd1 axis acts as a cholesterol sensor to decrease cellular accumulation of free cholesterol in order to diminish its harmful effects (44). Our immunohistochemical results showing similar protein localization of LXR and Srebp1 within the growth plate, as well as gene expression analysis showing similar expression patterns throughout chondrocyte differentiation, support this proposed mechanism.

Immunohistochemical analysis revealed LXR to be widely distributed in the embryonic growth plate, yet more localized to the pre-hypertrophic and resting zones upon activation with the LXR-specific agonist GW3965. Several studies in human macrophages and mouse adipose tissue have identified LXR response elements (LXRE) in the *Lxra* promoter, suggesting LXRs to possess tissue-specific autoregulatory functions (45–47). Our study suggests for the first time that such autoregulation exists in chondrocytes, where GW3965-treatment resulted in further preferential activation of LXRα in pre-hypertrophic and resting chondrocytes.

The overlap in LXR target genes between primary growth plate chondrocytes and IMACs was lower than expected. This could be due to cell type-specific effects, but changes in experimental design could also contribute to these differences. In particular, growth plate chondrocytes were incubated with GW3965 for 24 hours, while IMACs received the drug for 72 hours (19). Thus, secondary or tertiary effects of LXR activation would only be seen in the IMAC system. Nevertheless, the dominant biological process affected by LXR activation in both contexts was lipid metabolism, and the genes showing strongest induction overlapped greatly.

In summary, our study shows for the first time that LXR activation by its specific synthetic agonist, GW3965, differentially regulates genes involved in lipid metabolism in growth plate chondrocytes. Specifically, the LXR-Srebp1-Scd1 axis was identified to be a potential mechanism employed by LXR to maintain cellular cholesterol homeostasis. Further studies are needed to functionally validate these lipid metabolic changes in response to LXR activation in both the normal and diseased state. Better understanding of LXR’s role in this context will help shed light on the link between lipid homeostasis and the maintenance of chondrocyte health.

## Acknowledgements

The authors would like to thank all members of the Beier lab for ongoing support and discussions.

## Author Contributions

M.M-G.S. performed all experiments, data collection and statistical analysis. M.M-G.S. and F.B. participated in study conception, data interpretation, and manuscript drafting and editing. Both authors take responsibility for the integrity of this work.

