## Supplementary Tables for "Liver X Receptor activation regulates genes involved in lipid homeostasis in developing chondrocytes"

| Gene | Primer Sequence (5' – 3') |
|---|---|
| ***Acsl3*** | F - GAGGTCCAGCCATTGTTCAT<br>R - GTAATGATATGCCGCAGACG |
| ***Abca1*** | F - GCTACCCACCCTACGAACAA<br>R - GGAGTTGGATAACGGAAGCA |
| ***Scd1*** | F - CCCTCCTGCAAGCTCTACAC<br>R - CAGCCGTGCCTTGTAAGTTC |
| ***Abcg1*** | F - GCTGGGAAGTCCACACTCAT<br>R - CCTGCATGATGTAGCAGGAG |
| ***Srebf1*** | F - GATGTGCGAACTGGACACAG<br>R - CTGTCTCACCCCCAGCATAG |
| ***Lpcat3*** | F - CTACCCGTTGGCTCTGTTTT<br>R - AGCACGACACATAGCAAGGA |
| ***Actb*** | F - CTGTCGAGTCGCGTCCACCC<br>R - ACATGCCGGAGCCGTTGTCG |
| ***Gapdh*** | F - GCACAGTCAAGGCCGAGAATR<br>R- GCCTTCTCCATGGTGGTGAA |

Supplementary Table 1. Primer sequences used for RT-qPCR.

| Gene | Fold Change |
|---|---|
| *Abcg1* | 5.68304 |
| *Abca1* | 3.66906 |
| *Rdh13* | 2.83959 |
| *Srebf1* | 2.64691 |
| *Mir466g* | 2.04943 |
| *Mylip* | 2.04233 |
| *Lpcat3* | 1.85742 |
| *Apoc1* | 1.70349 |
| *Vmn1r36* | 1.70215 |
| *Ahcy* | 1.68003 |
| *Mir30e* | 1.67372 |
| *Fut11* | 1.66427 |
| *Scd1* | 1.63351 |
| *Gramd1b* | 1.58205 |
| *Eif2ak2* | 1.58131 |
| *Amy2b* | 1.57051 |
| *n-R5s218* | 1.55454 |
| *Gm7367* | 1.55335 |
| *Tcra-V22.1* | 1.55143 |
| *Scd2* | 1.53874 |
| *Acsl3* | 1.53432 |
| *Gm14444* | 1.51726 |
| *4930578G10Rik* | 1.50541 |
| *mmu-mir-466c-1* | -1.51245 |
| *Gm12888* | -1.56231 |
| *Olfr1336* | -1.67054 |
| *Vmn2r95* | -1.73502 |
| *Hist1h3a* | -1.75174 |
| *Mir669h* | -1.77484 |
| *Traj7* | -1.89333 |

Supplementary Table 2. Complete list of differentially expressed genes with LXR activation.
